# *Aedes aegypti* vision-guided target recognition requires two redundant rhodopsins

**DOI:** 10.1101/2020.07.01.182899

**Authors:** Yinpeng Zhan, Diego Alonso San Alberto, Claire Rusch, Jeffrey A. Riffell, Craig Montell

**Affiliations:** Department of Molecular, Cellular, and Developmental Biology and the Neuroscience Research Institute, University of California, Santa Barbara, California 93106, USA; Department of Biology, University of Washington, Seattle, WA 98195, USA

**Keywords:** Host recognition, mosquito, wind tunnel, vision, rhodopsin, optomotor

## Abstract

Blood-feeding insects, such as the mosquito, *Aedes* (*Ae*.) *aegypti*, use multiple senses to seek out and bite humans [1, 2]. Upon exposure to CO_2_, the attention of female mosquitoes to potential human targets is greatly increased. Female mosquitoes use vision to assist them in honing in on hosts that may be up to 10 meters away [3–9]. Only after coming into close range do convective heat from skin and odors from volatile organic compounds come into play, allowing female mosquitoes to evaluate whether the object of interest might be a host [10, 11]. Here, using CRISPR/Cas9 we mutated the gene encoding Op1, which is the most abundant of the five rhodopsins expressed in the compound eyes of *Ae. aegypti*. Using a cage assay and a wind tunnel assay, we surprisingly found that elimination of *op1* did not impair CO_2_-induced target seeking. We then mutated *op2*, which encodes the rhodopsin most similar to Op1, and also found that there was no impact on this behavior. Rather, mutation of both *op1* and *op2* was required to abolish vision-guided target recognition. In contrast to this defect, the double mutants still exhibited normal light attraction. By measuring the optomotor response, we found that the double mutants still recognized moving cues in their environment. In further support of the conclusion that the double mutant is not blind, we found that the animals retained an electrophysiological response to light, although it was diminished. This represents the first perturbation of vision in mosquitoes and indicates that host-seeking by *Ae. aegypti* depends on redundant rhodopsins.

## Results and Discussion

The mosquito, *Aedes* (*Ae*.) *aegypti*, infects ~80,000,000 people each year with flaviviruses that cause diseases ranging from dengue to yellow fever, chikungunya, West Nile, and Zika [12]. Only females bite, and they do so because they require nutrients from blood meals for egg development [13]. Unlike *Anopheline* malaria vectors, which bite primarily at night*, Ae. aegypti* seek out humans during daylight, particularly around dawn and dusk [14, 15]. Upon detecting CO_2_ plumes from human breath, female *Ae. aegypti* become much more responsive to visual cues and seek out hosts from ranges of up to several meters [3–8]. *Ae. aegypti* are especially attracted to people wearing dark clothing [16–18]. Even in the absence of humans, CO_2_ stimulates mosquitoes to seek out darker over lighter images [17, 18]. As a consequence of their poor visual acuity [19, 20], a dark spot is sufficient to attract them [3]. Once they are visually guided to within a few centimeters of the potential target, thermal and olfactory stimuli are the most salient in allowing the female mosquitoes to determine whether the image is an actual human, and these latter sensations are also enhanced by CO_2_ detection [2].

Despite the importance of the integration of CO_2_ and visual stimuli for long-range host detection in diurnal mosquitoes, there are no studies dissecting the roles of rhodopsins or other signaling proteins required for vision in any insect disease vector. The *Aedes* genome encodes 10 opsins, although only 5 are expressed in the compound eye [21]. To uncover the molecular mechanisms of vision-guided target recognition, we set out to explore the potential role of GPROp1 (Op1), since this long-wavelength visual pigment is the most widely-expressed rhodopsin in the eyes. There are eight photoreceptor cells (R1-R8) in each repeat unit (ommatidium) in the compound eye, and Op1 is expressed in the six outer photoreceptor cells (R1-6) and most R8 cells [22].

We used CRISPR/Cas9 genome editing to generate two independent *op1* alleles via homology-dependent repair. The *op1^R^* allele includes a 10 base-pair deletion and an insertion of the *DsRed* gene after the sequence encoding residue 72, near the second transmembrane domain (Figure 1A). The *op1^G^* mutation is characterized by an 8 base pair deletion and an insertion of the *GFP* gene after residue 82 (Figure 1A). We generated homozygous lines, which we confirmed by PCR (Figure S1A) and DNA sequencing. We performed real-time quantitative PCR and found that the *op1* RNA was dramatically reduced in both alleles (Figure 1B). We generated antibodies against Op1, and confirmed that Op1 protein was undetectable (Figure 1C).

**Figure 1.**
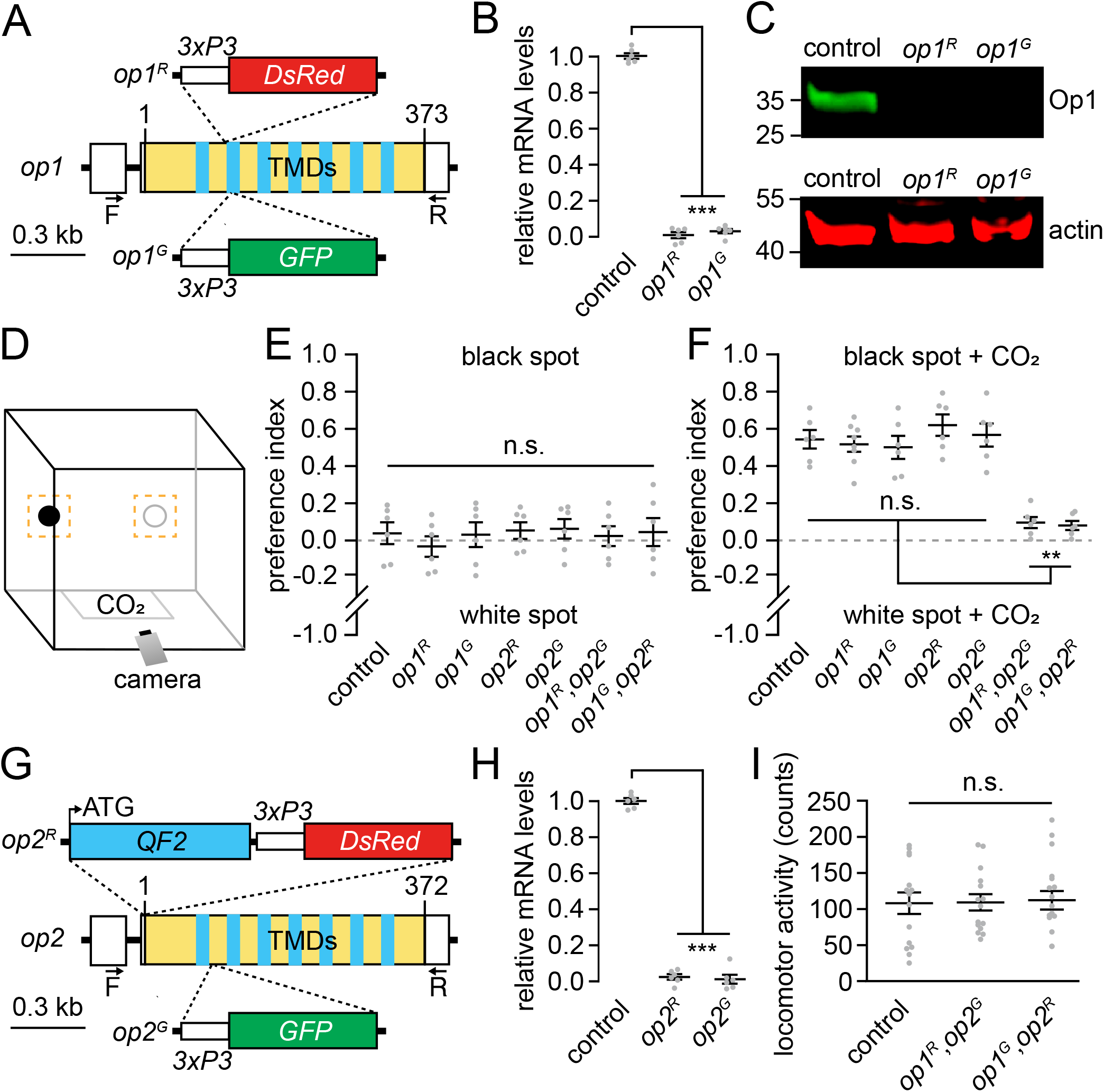
Generation of *op1* and *op2* mutants and cage assay for CO_2_-induced target selection. (A) Genomic structure of the *op1* gene and illustration of *op1^R^* and *op1^G^* alleles generated by CRISPR/Cas9-mediated HDR. (B) Normalized mRNA expression levels of *op1* alleles compared to the wild-type control determined by quantitative real-time PCR (n=6). (C) Western blot containing extracts from wild-type control, *op1^R^*, and *op1^G^* heads probed with anti-Op1 and anti-actin. The positions of protein size markers (in kilodaltons) are shown to the left. (D) Schematic of the cage for vision-guided host-seeking assay. The orange squares around the black and white circles represent the regions used to document the trajectories. (E) Vision-guided target selection cage assay showing the preference indexes of the indicated female mosquitoes for the black versus the white spot under clean air conditions. n=6 (50 females/assay). Total trajectories: control, n=23—51; *op1^R^*, n=35— 64; *op1^G^*, n=17—86; *op2^R^*, n=29—51; *op2^G^*, n=30—64; *op1^R^,op2^G^*, n=24—48; *op1^G^,op2^R^*, n=19—42. (F) Vision-guided target selection cage assay showing the preference indexes of the indicated female mosquitoes for the black versus the white spot exposed to 5% CO_2_. n=6 (50 females/assay). Total trajectories: control, n=87—146; *op1^R^*, n=52—183; *op1^G^*, n=45—167; *op2^R^*, n=63—129; *op2^G^*, n=76—204; *op1^R^,op2^G^*, n=34—51; *op1^G^,op2^R^*, n=26—60. (G) Genomic structure of the *op2* gene and the *op2^R^* and *op2^G^* alleles generated by CRISPR/Cas9-mediated HDR. (H) Normalized mRNA expression level of *op2* alleles compared to the wild-type control determined by quantitative real-time PCR (n=6). (I) Locomotor activities monitored over a 24-hour period in a DAM system. The counts are the number of times the animals broke the infrared beam. n=15. One-way ANOVA with Tukey’s multiple comparisons test for (B) and (H), and Kruskal-Wallis test with Dunn’s multiple comparison test for (E, F, and I). Means ±SEMs. Mann-Whitney U test with Bonferroni correction. n.s., not significant. **P<0.01. ***P<0.001.

To test whether mutation of *op1* impairs CO_2_-stimulated host target attraction, we devised an assay in a modified 30 x 30 x 30 cm insect cage that is based conceptually on work using wind tunnels [3]. The previous wind tunnel study showed that in the presence of a stream of CO_2_, female mosquitoes are attracted to a black feature, which serves as a surrogate host [3]. To create a simplified assay for measuring CO_2_-stimulated visual guidance to a surrogate host, we placed two small solid black and white circles at one end of a cage (Figure 1D). We inserted 50 females that were not blood fed into the cage and recorded their trajectories for 30 minutes (no added CO_2_).

We then introduced a source of 5% CO_2_, which increased their activity, and recorded their trajectories for 3 minutes. We quantified the total number of times the mosquitoes explored a small fictive area (6 cm x 6 cm) around the black and white circles and calculated a preference index.

We found that under standard air conditions control females had no preference for either the black versus the white spot (Figure 1E), or for the black spot versus another random area in the cage (Figure S2). In contrast, upon exposure to CO_2_ the control females exhibited a strong preference for the black spot (Figure 1F), reminiscent of the previous experiments using a wind tunnel [3]. We then analyzed the *op1^R^* and *op1^G^* mutants. In the absence of CO_2_ stimulation, they had no bias for either the black or white spots similar to the response seen in controls (Figure 1E). Upon addition of CO_2_, the *op1* mutant females were as attracted to the dark feature as control animals (Figure 1F). This was unexpected since Op1 is the major rhodopsin in the compound eyes [22]. These data indicate that Op1 is either irrelevant for visual recognition of a surrogate host or is functionally redundant with another rhodopsin.

The rhodopsin most related to Op1 is GPROp2 (Op2), which is 89.8% identical to Op1 and is expressed in a subset of R7 photoreceptor cells [21, 23]. The next most related visual rhodopsins, Op3 and Op7, are 79.9 and 64.3% identical, respectively. Op2 is also a long-wavelength rhodopsin, similar to Op1 [21]. We used CRISPR/Cas9 to generate two *op2* alleles. *op2^R^* has a 14 base pair deletion and an insertion of the *DsRed*gene at the start codon, and *op2^G^* has an 11 base pair deletion beginning after the sequence encoding residue 75 and an insertion of the *GFP* gene (Figure 1G). We verified the mutations by PCR (Figure S1B) and DNA sequencing and confirmed that expression was eliminated by real-time quantitative PCR (Figure 1H). Using the cage assays, we tested CO_2_-stimulated visual target attraction and found that the *op2* mutants behaved similarly to the controls (Figure 1F).

Due to the lack of impairment resulting from mutation of either *op1* or *op2* alone, we generated and tested the effects of two double mutants: *op1^R^,op2^G^* and *op1^G^,op2^R^*. The double mutant females exhibited normal locomotor activity using a TriKinetics DAM assay system, which counts the number of times animals cross an infrared beam over a 24-hour period (Figure 1I). We then tested the attraction of the *op1,op2* double mutants to the black feature in the cage assay. Under standard air conditions, the double mutant females had no preference for the black versus the white circles, as is the case for the controls (Figure 1E). However, both *op1,op2* alleles exhibited a dramatic deficit in CO_2_-stimulated visual target attraction (Figure 1F).

In a natural environment detection of a CO_2_ plume induces female *Ae. aegypti* to surge upwind and use vision to find a human host [3, 5–7]. This behavior has been modeled in the laboratory using a wind tunnel [3]. To determine whether the phenotypes exhibited using the cage assay reflected their behavior in the presence of wind, we employed an assay similar to that previously described [3]. We used a large wind tunnel that spans 400-body lengths of the mosquito (equivalent to 8 football fields for a human), which contained identically sized black and white spots. We inserted 50 nonblood fed female mosquitoes into the apparatus and recorded flight trajectories using a three-dimensional tracking system that includes 16 cameras (Figure 2A). When control females are exposed to air with normal, atmospheric levels of CO_2_ (~400 ppm) they fly randomly and there is no preference for the black spot over the white spot, or in comparison to elsewhere in the tunnel (Figure 2B and 2H). As previously described [3], after introducing a 10% CO_2_ plume into the tunnel (~100,000 ppm) control females greatly increase their exploration of the dark black object (Figure 2C). This change is large as the percent of trajectories in a 14 x 14 x 4 cm vicinity of the black spot increased more than 5-fold in the presence of CO_2_ versus standard air (11.7% vs. 2.3%; Figures 2H and 2I). We found that the *op1^R^* and the *op2^R^* single mutant females exhibited behaviors indistinguishable from the controls. Under standard air conditions they were indifferent to the black spot that represented a surrogate host (Figures 2D, 2E and 2H), and upon exposure to 10% CO_2_ they showed the same attraction to the black feature and overall flight trajectories as control females (Figures 2F, 2G, 2I—2K).

**Figure 2.**
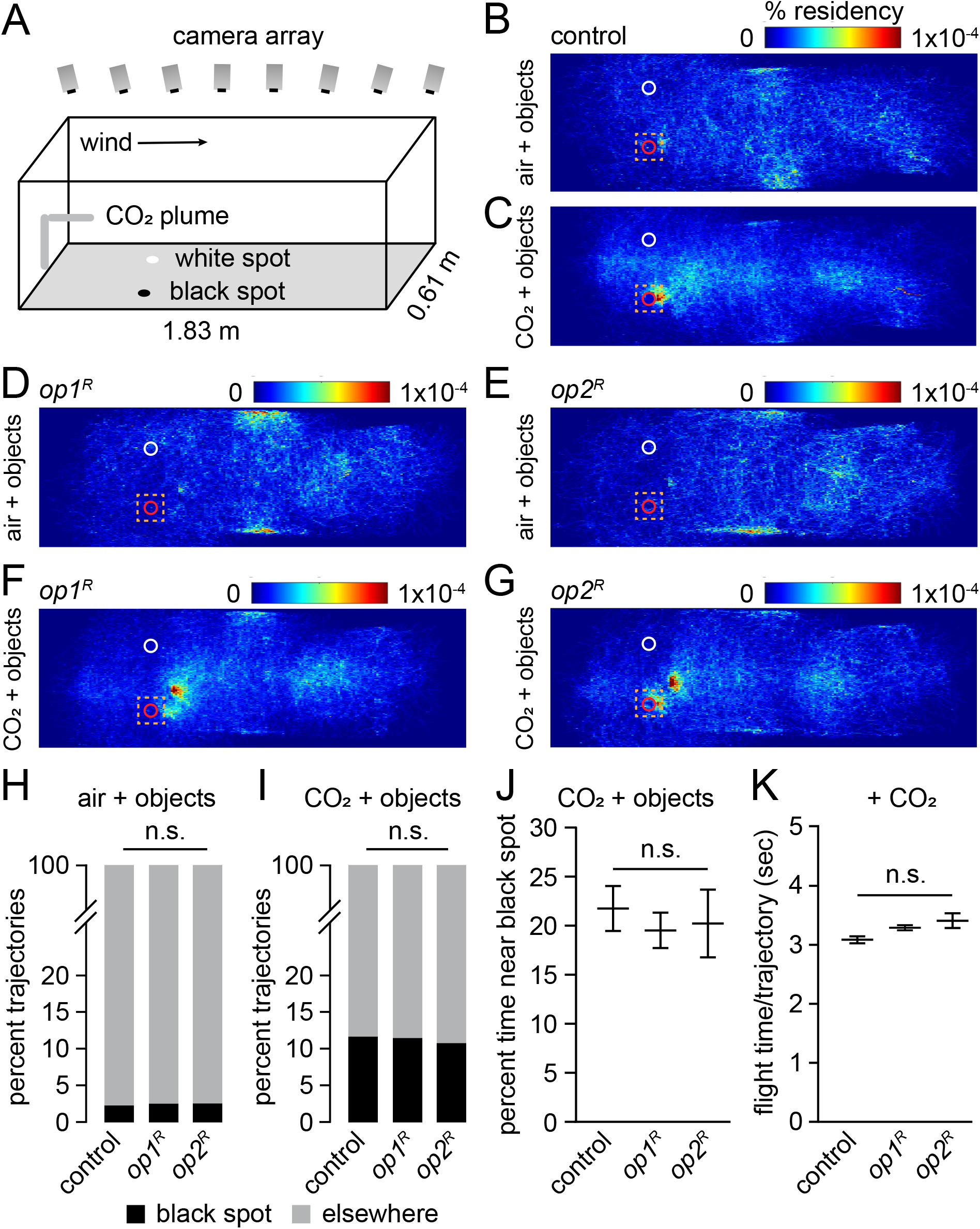
Wind tunnel assays showing *op1* and *op2* single mutants have normal vision-guided target-recognition. (A) Schematic of the mosquito wind tunnel assay with the 3D-tracking system. (B—G) Heat-maps showing the flight trajectories of the indicated females exposed to standard air or 10% CO_2_. The red and white circles indicate the black and white spots, respectively. The orange squares define the regions used to document the trajectories near the black or white objects. The colored scale bars indicate the % residency. (B) Control exposed to standard air. (C) Control exposed to 10% CO_2_. (D) *op1^R^* exposed to standard air. (E) *op2^R^* exposed to standard air. (F) *op1^R^* exposed to 10% CO_2_. (G) *op2^R^* exposed to 10% CO_2_. (H) Percent of detected trajectories near the black spot versus other places under standard air conditions. Total trajectories: control, n=1,567; *op1^R^*, n=5,107; *op2^R^*, n=1,191. (I) Percent of detected trajectories near the black spot versus other places when exposed to 10% CO_2_. Total trajectories: control, n=4,689; *op1^R^*, n=11,438; *op2^R^*, n=1,442. (J) Percent of time that the females spent near the black spot versus other places in the presence of 10% CO_2_. Total trajectories: control, n=387; *op1^R^*, n=581; *op2^R^*, n=130. (K) Average flight times per trajectory exhibited by the indicated females exposed to 10% CO_2_. Total trajectories, control, n=4,689; *op1^R^*, n=11,438; *op2^R^*, n=1,442. Fisher’s exact test for (H) and (I), and Mann-Whitney U test with Bonferroni correction. Bootstrapped 95% confidence interval of the mean for (J) and (K). n.s., not significant.

We then tested the double opsin mutants in the wind tunnel. In the absence of added CO_2_, the *op1^R^,op2^G^* and *op1^G^,op2^R^* females were not attracted to the black spot, similar to results seen in the controls (Figures 3A, 3B and 3E). However, upon exposure to CO_2_, the double mutants did not exhibit any elevation in trajectories in the vicinity of the black feature relative to their behavior in standard air (Figures 3C, 3D and 3F). In addition, the percentage of time that they spent near the black spot was greatly reduced compared to the control females (Figure 3G). Importantly, their flight time per trajectory was similar to controls (Figure 3H), indicating that the defect in object recognition is not attributable to loss of sustained flight. Overall, these data indicate that the cage assay can provide a simplified alternative to a wind tunnel assay in laboratories that do not have access to such wind tunnels. Moreover, the findings with the cage assay and wind tunnel support the conclusion that Op1 and Op2 have redundant roles necessary for CO_2_-induced vision-guided target attraction.

**Figure 3.**
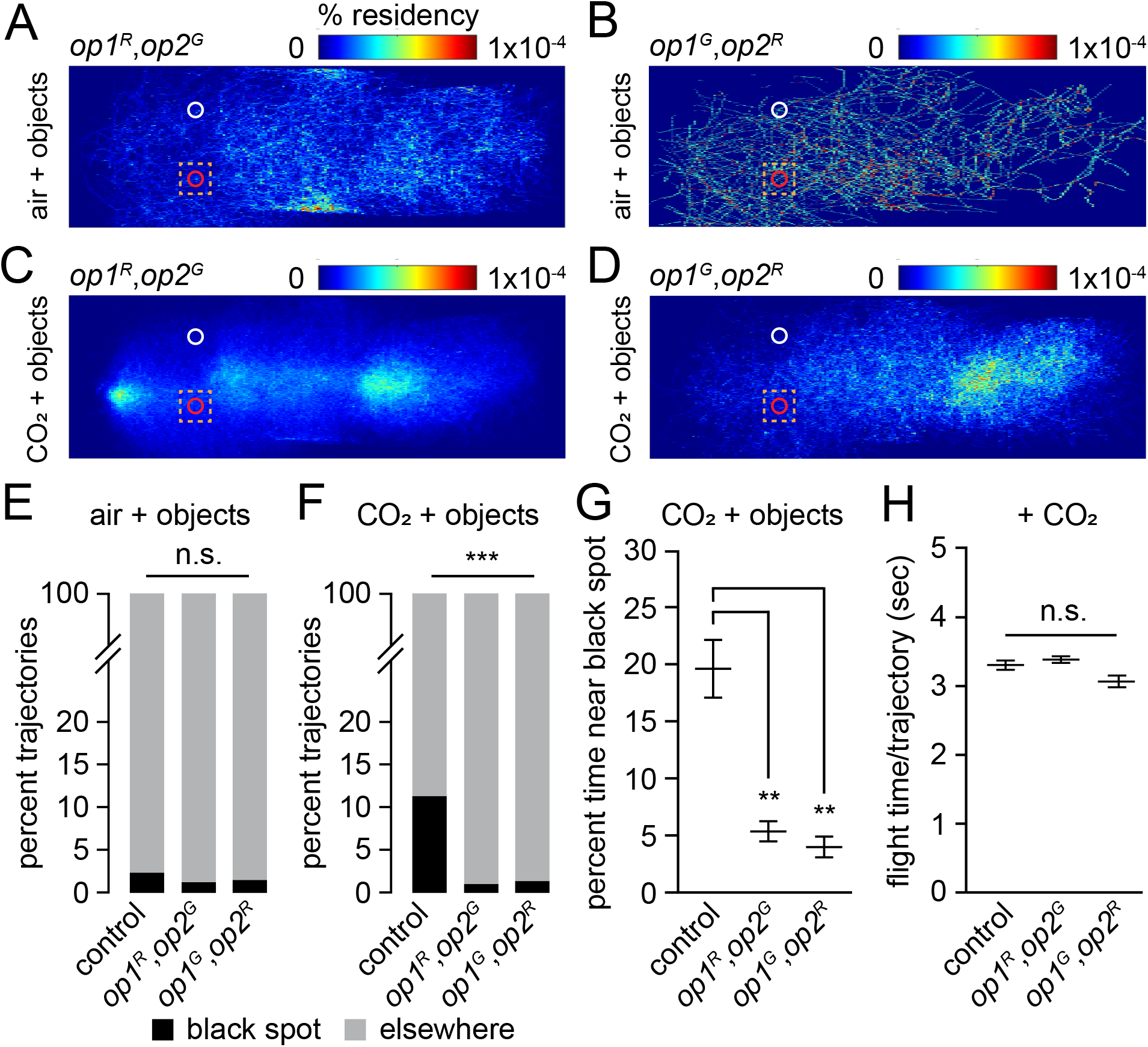
Elimination of vision-guided target-recognition in *op1,op2* double mutants. (A—D) Heat-maps showing the flight trajectories of *op1,op2* double mutant females exposed to standard air or 10% CO_2_. The red and white circles indicate the black and white spots. The orange squares define the regions used to document the trajectories near the black or white objects. The colored scales bar indicate the % residency. (A) *op1^R^,op2^G^* exposed to standard air. (B) *op1^G^,op2^R^* exposed to standard air. (C) *op1^R^,op2^G^* exposed to 10% CO_2_. (D) *op1^G^,op2^R^* exposed to 10% CO_2_. (E) Percent of detected trajectories near the black spot versus other places under standard air conditions. Total trajectories: control, n=2,229; *op1^R^,op2^G^*, n=1,624; *op1^G^,op2^G^*, n=405. (F) Percent of detected trajectories near the black spot versus other places when exposed to 10% CO_2_. Total trajectories: control, n=4,802; *op1^R^,op2^G^*, n=8,883; *op1^G^,op2^R^*, n=2,250. (G) Percent of time mosquitoes spent near the black spot versus other places in the presence of 10% CO_2_. n=86—5581 total trajectories. Total trajectories: control, n=289; *op1^R^,op2^G^*, n=91; *op1^G^,op2^R^*, n=66. (H) Average flight time per trajectory exhibited by the indicated mosquitoes in the presence of 10% CO_2_ condition. Total trajectories: control, n=4,802; *op1^R^,op2^G^*, n=8,883; *op1^G^,op2^R^*, n=2,250. Fisher’s exact test for (E) and (F), and Mann-Whitney U test with Bonferroni correction. Bootstrapped 95% confidence interval of the mean for (G) and (I). Kruskal-Wallis test with Dunn’s multiple comparisons test for (H), Means ±SEMs. n.s., not significant. **P<0.01, ***P<0.001.

The inability of the *op1,op2* double mutant mosquitoes to recognize the visual target (surrogate host), raised the possibility that the animals were unable to detect visual stimuli or even light. To determine whether the mutant mosquitoes were capable of responding behaviorally to light, we tested whether these animals exhibited positive phototaxis upon stimulation with CO_2_. We found that the single and double mutants all exhibited phototaxis indexes similar to the controls (Figure 4A). Therefore, the *op1,op2* double mutants retain the ability to sense ambient light.

**Figure 4.**
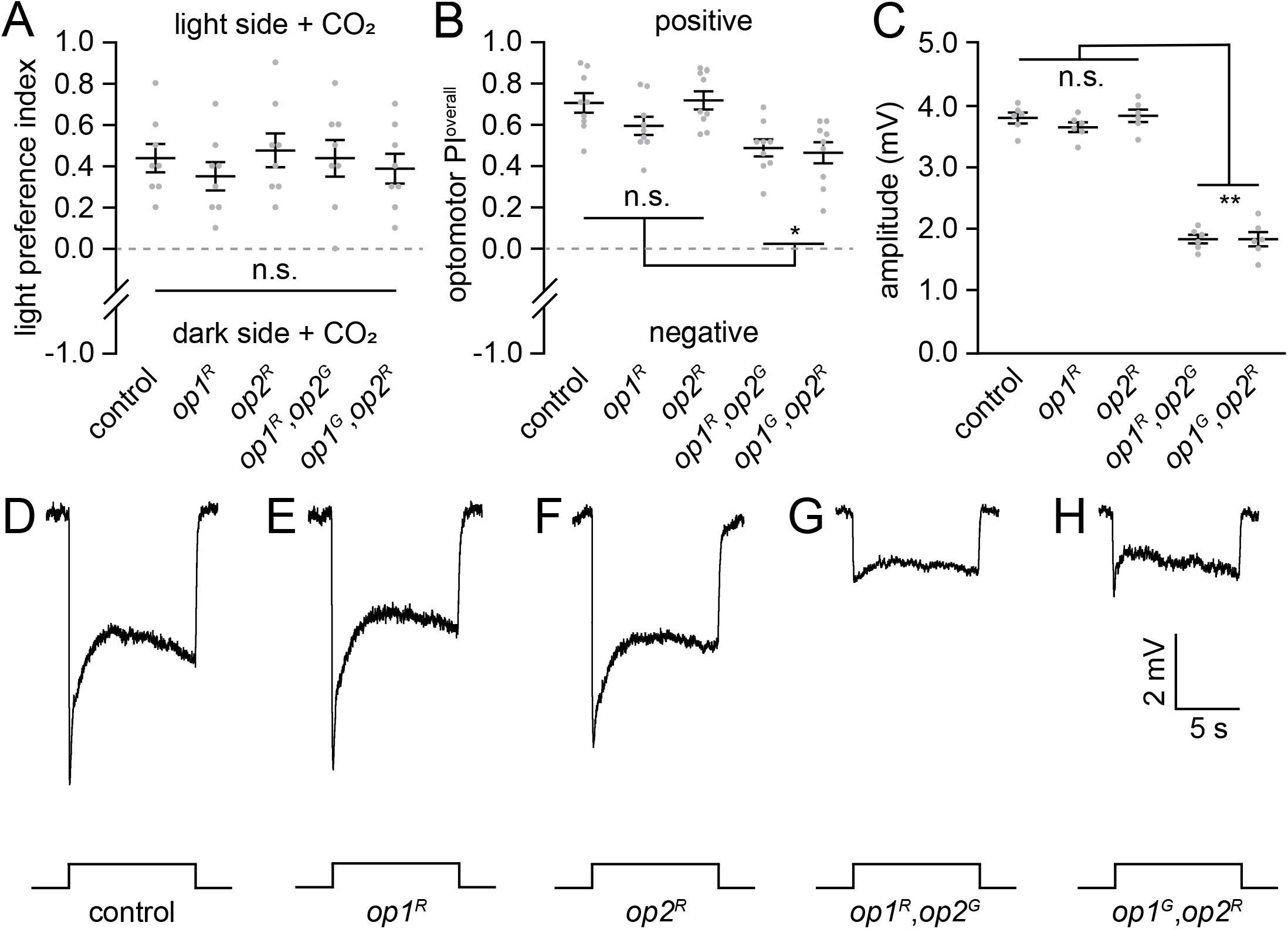
Phototaxis, optomotor responses and ERG responses exhibited by *opsin* mutants. All assays were performed using females. (A) Phototaxis assays exhibited by the indicated animals. n=8 (30 females/assay). (B) Optomotor response exhibited by the indicated animals. n=9. (C) ERG amplitudes exhibited by the indicated animals. n=6. (D—H) ERG recordings in response to a 10 sec white light pulse (~1000 lux). (D) Control. (E) *op1^R^*. (F) *op2^R^*. (G) *op1^R^,op2^G^*. (H) *op1^G^,op2^R^*. Statistics performed using the Kruskal-Wallis test with the Dunn’s multiple comparisons test for (A) and (B). ANOVA with Tukey’s multiple comparisons test for (C). Means ±SEMs. n.s., not significant. **P<0.01, *P<0.05.

To ascertain whether or not the mutants were capable of recognizing moving objects we examined the optomotor response, which is an innate orienting behavior evoked when objects in the surrounding environment are moving [24]. We placed a single female mosquito in the center of a rotating drum with alternating black and white vertical stripes. This causes wild-type mosquitoes to walk in the same direction as the moving stripes [25]. Both *op1,op2* double mutants favored walking with the moving black stripes, although the response was reduced (Figure 4B). These findings indicate that the *op1,op2* double mutant mosquitoes were capable of seeing and tracking moving objects, even though their capacity to recognize the surrogate host in the wind tunnel assay was virtually eliminated. Thus, on the basis of the phototaxis and optomotor responses, we conclude that the double mutants are not blind.

To assess the visual response further, we performed electroretinogram (ERG) recordings, which are extracellular recordings that measure the summed responses of all retinal cells to light. Upon initiation of a light stimulus, control mosquitoes exhibit a corneal negative response, which quickly declines to the baseline upon cessation of the light stimulus (Figures 4C and 4D). The ERG responses of the *op1^R^* and *op2^R^* single mutants were indistinguishable from the control (Figures 4C, 4E and 4F). The double mutant mosquitoes also exhibited ERG responses, although they were reduced relative to the control (Figures 4C, 4G and 4H). These findings are consistent with the optomotor results demonstrating that the double mutants are visually impaired, but not blind.

In *Drosophila*, motion detection is primarily mediated by R1-6 photoreceptor cells [26, 27]. *Ae. aegypti op1* is expressed in R1-6 photoreceptor cells and most R8 photoreceptor cells [22], so we expected loss of Op1 to have a profound effect on vision-guided target attraction. The surprising finding that mutation of *op1* had no impact on this behavior indicated that Op1 is either irrelevant for honing in on targets, or that another rhodopsin is required. We knocked out *op2* since it encodes the rhodopsin most related to Op1, among the four other rhodopsins expressed in the compound eye. However, mutation of *op2*, which is expressed in R7 photoreceptor cells [21, 23], also had no effect on CO_2_-induced target recognition.

Our finding that target recognition is eliminated only upon mutation of both *op1* and *op2* suggests that they function redundantly with respect to host identification. In dipterans, the R1-6 photoreceptor cells send their axons to the lamina in the optic lobe, while the R7 and R8 photoreceptor cells extend their axons through the lamina to specific layers in the next region in the optic lobe, the medulla [24, 28]. Although activation of Op1 and Op2 leads to stimulation of distinct neuronal pathways, we suggest that the different pathways ultimately converge onto the same region of the central brain that functions in the detection of a static object during flight. Consequently, activation of either Op1 or Op2 is sufficient. Finally, the discovery that mutation of *op1* and *op2* virtually eliminates visual guidance to a potential host raises the possibility that using gene drive approaches [29–32] to mutate these genes in mosquito populations would interfere with their visual attraction to humans and reduce the incidence of diseases, such as dengue, which affect millions of people.

## Acknowledgments

This work was supported by a grant to C.M. from the NEI and by a grant to J.A.R from the NIAID. We thank D. Thakur for comments on this manuscript and N. DeBeaubien for comments on this manuscript and for preparation of the cage for performing the target attraction assay. We thank O.S. Akbari for the *Ae. aegypti* (Liverpool strain) and for the AAEL006511-Cas9 transgenic *Ae. aegypti* line expressing *Cas9* under control of the *ubiquitin L40* promotor. We also thank M. Li and O.S. Akbari for advice enabling us to improve our embryo injections. C.M. also thanks A.A. James for early discussions that motivated C.M. to initiate work on mosquitoes.

## Author contributions

Y.Z. and C.M. designed the experimental plan, analyzed the data, and prepared the manuscript. C.M. also supervised the research and obtained the funding for this study. Y.Z. conducted most of the assays, except for the wind tunnel experiments, which were conducted by D.A.S.A., C.R. and J.A.R. D.A.S.A., C.R. and J.A.R. analyzed the data, and improved the manuscript with their comments.

## Competing Financial Interests

The authors declare no competing financial interests.

## Methods

### Mosquito rearing

The control mosquitoes used were *Ae. aegypti*(sequenced Liverpool strain; gift from O.S. Akbari), which were reared at 28°C, 80% relative humidity under a 14hr:10hr light:dark regime in walk-in chambers, located in an ACL-2 facility. Mosquito eggs were hatched in deionized water and fed fish food (TetraMin tropical granules, Tetra) until the emergence of pupae. Male and female mosquitoes were sorted at the pupal stages based on their sizes and then transferred into an insect collection cage (17.5 x 17.5 x 17.5 cm, BugDorm) for mating and maintaining. Adult mosquitoes were fed 10% sucrose placed on cotton balls. To promote egg production, adult females (5—10 days old) were fed blood using an artificial feeder (Hemotek) heated with fresh defibrinated sheep blood (Hemostat). All mutant mosquitoes were generated using the Liverpool line of *Ae. aegypti* and outcrossed to this background for eight generations.

### Generation of transgenic strains

To generate the *op1* and *op2* alleles, we selected short-guide RNAs (sgRNAs) that targeted the *GPROp1* (LOC5568060) and *GPROp2* (LOC5567680) loci using the CRISPR Optimal Target Finder (https://flycrispr.org/target-finder/). The target sequences of the sgRNAs used for generating the *op1* and *op2* alleles are presented in Table S1.

To generate the *op1^R^, op1^G^* and *op2^G^* alleles, we created the *op1^R^-3xP3-DsRed-HDR, op1^G^-3xP3-GFP-HDR* and *op2^G^-3xP3-GFP-HDR* DNA constructs for microinjections. To do so, we used the In-Fusion cloning kit (Clontech) to introduce the sgRNAs, and the upstream and downstream homology arms (~1 kb each) into the pAeU6-LgRNA-3xP3-DsRed or the pAeU6-LgRNA-3xP3-GFP vectors (J. Chen, J. Luo, and C.M., unpublished), which includes the sequences encoding the 3xP3-driven fluorescent marker, SV40 transcription terminator, the *U6* promoter for directing expression of the sgRNA, and the gRNA scaffold. In addition to the insertion of the genes encoding either DsRed or GFP, *op1^R^* has a 10 base pair deletion, which begins after the sequence encoding residue 72, *op1^G^* has an 8 base pair deletion, which begins after the sequence encoding residue 83, and *op2^G^* has an 11 base pair deletion, which begins after the sequence encoding residue 75.

The *op2^R^* allele includes a 14 base pair deletion, which removes the ATG and the following 11 base pairs. To create this allele, we first generated the pU6-gRNA1 plasmid (Figure S3), which includes the sequences encoding QF2, the 3xP3-driven fluorescent marker, SV40 transcription terminator, the *U6* promoter for directing expression of the sgRNA, and a gRNA scaffold. QF2 was cloned from pBAC-DsRed-QF2-hsp70 (Addgene #104876). We then used pU6-gRNA1 to introduce the sgRNAs, and the upstream, and downstream homology arms (~1 kb each). The primers used for cloning the homology arms are presented in Table S2.

All transgenic strains were generated by microinjecting the plasmids into embryos of the transgenic *Ae. aegypti* line that expresses Cas9 under control of the *ubiquitin L40* promotor (gift from O.S. Akbari) [33]. Briefly, we collected freshly laid embryos, and microinjected the plasmid DNA (~500 ng/μL) into the posterior ends of ~2000 embryos using a micro-injector (Eppendorf) and a Zeiss Axioplan 2 microscope. G0 embryos hatched four days post-injection, and the adult G0 animals (~100 per injection) were crossed to the opposite sex. The females were blood-fed to generate G1 progeny. The eyes of the G1 larvae were screened for expression of the DsRed or GPF fluorescent markers under a Zeiss SteREO Discovery. V8 stereomicroscope. Positive G1 animals were genotyped by PCR and out-crossed to the wild-type control strain for eight generations. We genotyped the homozygous lines before performing experiments, using the primers listed in Table S3.

### Generation of Op1 antibodies and Western Blots

Rat anti-Op1 was generated (Pocono Rabbit Farm & Laboratory, Inc.) to a peptide that spanned amino acid residues 338 to 356 (<u>CTQKFPALSSTDAPAASNSD</u>). To perform the protein blots, 20 heads from 5-7 day-old female mosquitoes were homogenized, fractioned by SDS-PAGE (Bio-Rad Mini-Protean TGX Gels, 4-15%, Cat # 456-1086), transferred to nitrocellulose membranes (Bio-Rad, cat. # 162-0112), probed with anti-Op1 (1:20 dilution) and rabbit anti-actin (Abcam, cat. # ab1801) as the loading control. The blots were then incubated with IRDye-conjugated goat anti-rat secondary antibodies (1:5000 dilution; LI-COR Biosciences, cat. #926-32219), and goat anti-rabbit secondary antibodies (1:5000 dilution; LI-COR Biosciences, cat.#926-68071), and visualized using a LI-COR imager system (Odyssey^®^ CLx).

### Quantitative PCR (RT-qPCR)

To detect the expression of *op1* and *op2* RNAs in the control and mutant mosquitoes, we extracted total RNA from 10 heads with Trizol reagent (Thermo Fisher Scientific). 1 μg RNA was reverse transcribed using Reverse Transcription Kit (Promega) with oligo(dT) primers. RT-qPCR was carried out using a LightCycler 480 SYBR Green I Master Mix (Roche). *Rpl32* (LOC5577996) was used as the normalization reference. The qPCR primers are listed in Table S4.

### Vision-guided target-attraction using a cage assay

The cages used in this assay were modified by N. DeBeaubien from a standard 30 x 30 x 30 cm cage (BugDorm). The mesh material on one of the vertical sides of the cage was replaced with a 12” x 12” x 1/16” clear cast acrylic sheet (McMaster-Carr; catalog ID: 8560K171) to allow for clear video recording and tracking. To conduct the vision-guided target-attraction assays, we glued a 3 cm black circle (generated by an HP, LaserJet Pro MFP M426fdn), and a 3 cm white circle to the interior wall of the cage so that they were separated by 18 cm. We moved the cage into a walk-in chamber held at 28° and 80% humidity. A CO_2_ Flypad (Genesee Scientific; cat. # 59-119) was placed below the cage near the front of the wall that had the visual cues. Before initiation of the test, the CO_2_ was kept in the off position.

To perform the assays, we used 4—10-day-old, non-blood-fed females that were sucrose deprived for 48 hours, but had access to water. The animals were maintained under 14-hr light:12-hr dark cycles. We inserted 50 females into the cage and the assays were initiated 2 hours before lights off (ZT12). We then recorded their trajectories on the wall that had the visual cues for 30 minutes (standard air condition) with a webcam (Logitech C922) at 30 frames/sec. We then introduced 5% CO_2_ through the Flypad and recorded the mosquitoes’ trajectories on the wall with the visual cues for 3 min (CO_2_ condition). The preference index (PI) was calculated as follows: PI = (N^B^ – N^W^)/(NB + N^W^). NB = total number of times that the mosquitoes explored the black spot within the fictive area (6 x 6 cm). N^W^ = total number of times that mosquitoes exploring the white spot within the fictive area (6 x 6 cm), or a random area with the same size as white area (Figure S2B). A PI = 0 indicates no preference between the two spots. A positive PI (>0—1) indicates that the mosquitoes prefer the black spot, and a negative PI (−1 —<0) indicates that the mosquitoes prefer the white spot or the random area.

### Locomotor activity

Measurements of total locomotor activity over a 24-hr period were conducted at 28°C using the Drosophila Activity Monitoring (DAM) System (TriKinetics, LAM25). Individual 3—5 day-old non-blood-fed female mosquitoes were inserted into monitoring tubes (TriKinetics, PGT 25 x 125 mm Pyrex Glass), which contained 10% sucrose as a food source. Locomotor activity was determined automatically over the course of 24 hr (ZT0—ZT24; 14 hr light and 10 hr dark) by automatic tabulation of the number of times the animals broke the infrared beam. Data acquisition and analyses were performed using the DAMSystem (TriKinetics) and MATLAB.

### Vision-guided target-seeking using a wind tunnel assay

All experiments were performed in a low-speed wind tunnel (ELD Inc., Lake City, MN), with a working section of 183 x 61 x 61 cm and a constant laminar air flow of 0.4 m/sec. A low contrast checkerboard was projected on the floor of the arena and a low contrast grey horizon was projected on each side of the arena [34]. We used three rear projection screens (SpyeDark, Spye, LLC, Minneapolis, MN) and three short-throw projectors (LG PH450U, Englewood Cliffs, NJ) to project background images. A 3D realtime tracking system was used to track the mosquitoes’ trajectories [34]. 16 cameras (Basler AC640gm, Exton, PA) were mounted on top of the wind tunnel and recorded mosquito trajectories at 60 fps. All cameras had an opaque Infrared (IR) Optical Wratten Filter (Kodak 89B, Kodak, Rochester, NY) to mitigate the effect of light in the tracking. IR backlights (HK-F3528IR30-X, LedLightsWorld, Bellevue, WA) were installed below and to the sides of the wind tunnel to provide constant lighting. Finally, two circles (one black and one white) that were 4-cm diameter were used as visual cues (Color-Aid, Hudson Falls, NY). These visual cues were placed perpendicular to the airflow and 15 cm downwind from the odor source, and were separated by 25 cm. 50 female mosquitoes, starved for at least 12 hr, were released inside the wind tunnel. The circadian rhythm of all mosquitoes was calibrated so that the experiment started 3 hr before sunset. For a single trial, mosquito trajectories were recorded for a total duration of 5 hr: one hr of clean air (an acclimation phase), 2 hr of a CO_2_ plume (10%), followed by 2 hr of clean air.

The CO_2_ and purified air were automatically delivered using two mass flow controllers MC-200SCCM-D (Alicat Scientific, Tucson, AZ) that were controlled by a python script that allowed synchronizing odor and air delivery with the trajectory behaviors. In the acclimation and post-CO_2_ periods of the experiment, only purified air was released from the odor nozzle, whereas during CO_2_ delivery the plume consisted of 90% air and 10% CO_2_. The CO_2_ plume was calibrated using a CO_2_ analyzer (LI-COR Biosciences). Only trajectories that lasted for more than 1.5 seconds were analyzed (total number >59,000). To examine mosquito visual attraction behaviors, a fictive volume was created around the visual cues (area: 14 x 14 cm; height: 4 cm).

### Walking optomotor response

Mosquito walking optomotor responses were conducted as previously described with modifications using 4—6 day-old, non-blood-fed female mosquitoes [35]. The mosquitoes were anesthetized on ice, the wings were removed, and the animals were allowed to recover for 3 hr before performing the behavioral experiments at Zeitgeber time 12—14, which is last two hours of the light cycle, since the animals were maintained under a 14 hr light:10 hr dark regime. Individual, wingless mosquitoes were placed in a 30 mm diameter 28°C chamber in the middle of a rotating drum (diameter, 10 cm; height, 22.5 cm), which had alternating black and white vertical stripes. The angular width of each stripe was 18°, and the drum was rotated at 30 RPM. White LED lights (~500 lux) were illuminated surrounding the drum during the experiment. To conduct each trial, the mosquito was tested for the optomotor response with the drum rotating in the clockwise direction for 120 sec. The drum then stopped rotating for 30 sec, after which it started rotating in the counterclockwise direction for another 120 sec. We documented the turning and walking response to the rotating visual field by videotaping with a webcam (Logitech C922) at 30 frames/sec.

To calculate the overall optomotor performance index (PI), we first calculated the individual PIs when the drum rotated either in the clockwise direction (PI^clockwise^) and counterclockwise direction (PI_counterclockwise_). The performance index (PI) was calculated according to the following equation: PI = (N^same^ - N^different^)/(N^same^ + N^different^). N^same^ = number of times the mosquito walked across a quadrant line in the same direction as the rotating stripe, and N^different^ = number of times the mosquito walked against the direction as the rotating stripe. PI = 0 means that the mosquito does not track the stimulus. A positive PI (>0—1) indicates that the mosquito exhibits a positive response, which is turning and walk the same direction as the visual stimulus. A negative PI (−1 — <0) indicates that the mosquito turns against the visual stimulus. The overall optomotor PI of each trial is defined as PI^overall^=(PI^clockwise^ + PI^counterclockwise^)/2.

### Phototaxis assay

To measure phototaxis, we performed the assays at Zeitgeber time 12—14, using 4—6 day-old, non-blood-fed females. We anesthetized 30 females on ice, introduced them into a transparent plastic tube (length, 15.2 cm; diameter, 3.8 cm; Uline, cat. # S-12642), plugged the open end with a cotton ball, and kept the animals in the dark for 30 min so that they become dark-adapted and their locomotor activity recovered. We then used transparent tape to connect the tube with the mosquitoes and cotton plug to another tube covered with black paper. We then gently removed the cotton ball between the two test tubes, exposing a 1 cm hole between the two tubes. To stimulate the animals with CO_2_, we quickly placed the tubes containing the mosquitoes on a CO_2_ Flypad (Genesee Scientific; cat. # 59-119), so that the hole was directly pressed up against the pad.

To initiate the assay, we gently shook the tube for 3 sec to distribute the mosquitoes on both sides, then simultaneously turned on a ~500 lux LED light (exposing the side of the set up that was not covered with black paper). After 1 min, the apparatus was photographed, and the number of mosquitoes on the light side were counted. The light preference index (PI) was calculated as follows: PI = (N^L^ - N^D^)/(N^L^ + N^D^). N^L^ = number of mosquitoes located on the side with the light. N^D^ = number of mosquitoes situated on the dark side. A PI = 0 indicates no preference between the two sides. A PI (>0—1) and (−1 —<0) indicates preference for the light and dark sides, respectively.

### ERG recordings

ERG recordings were performed by fixing 4—6 day-old, non-blood-fed female mosquitoes to a coverslip with beeswax. The reference and recording glass electrodes (thin-wall glass capillaries; OD, 1.0 mm; length, 76 mm; World Precision Instruments, cat. # TW100F-3) were pulled using a micropipette puller (Sutter Instrument, p-97), and filled with Ringer’s solution (3 mM CaCl_2_, 182 mM KCl, 46 mM NaCl, 10 mM Tris pH 7.2). The reference electrode was placed on the thorax in a small drop of electrode cream (Parker, cat. # 17-05), and the recording electrode was placed in electrode cream on the surface of the compound eye. Mosquitoes were exposed to a 10-sec pulse of white light (~1000 lux) from a light source (Apex illuminator, Newport), though a light guide. The light-induced responses were amplified by using an IE-210 amplifier (Warner Instruments) and digitalized using a Powerlab 4/30 device (AD Instruments). Data were visualized and analyzed with LabChart 6 software (AD Instruments).

### Quantification and Statistical Analysis

Statistical analyses were performed using Prism 8 (GraphPad Software), MATLAB and SPSS Statistics 26 (IBM). For wind tunnel experiments, we calculated the mean and performed bootstrapping of the 95% confidence interval of the mean by re-sampling random trajectories 1000 times. We performed a non-parametric Mann-Whitney U test with Bonferroni correction to determine the statistically significant of two groups. We conducted Fisher’s exact test for determining statistical significance of the contingency tables. To determine statistical significance with multiple comparisons, we either conducted parametric tests using one-way ANOVA with Tukey’s multiple comparisons test, or non-parametric tests using the Kruskal-Wallis test with Dunn’s multiple comparison test. The data are displayed as means with 95% CI or SEMs as indicated in each legend.

## Supplementary Figure legends

**Figure S1.**
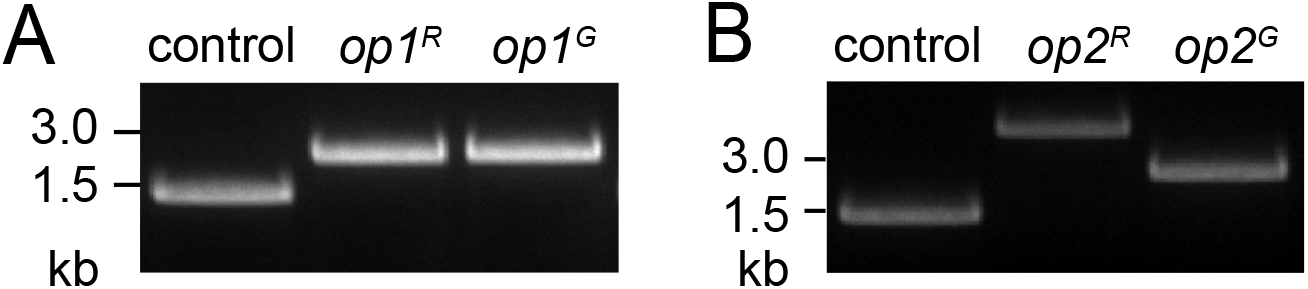
PCR genotyping of *opsin* mutations. (A) Genotyping the *op1^R^* and *op1^G^* mutations by PCR. The 5’ and 3’ primers are indicated by the arrows labeled F (forward) and R (reverse), respectively in Figure 1A. (B) Genotyping the *op2^R^* and *op2^G^* mutations by PCR. The 5’ and 3’ primers are indicated by the arrows labeled F (forward) and R (reverse), respectively in Figure 1G.

**Figure S2.**
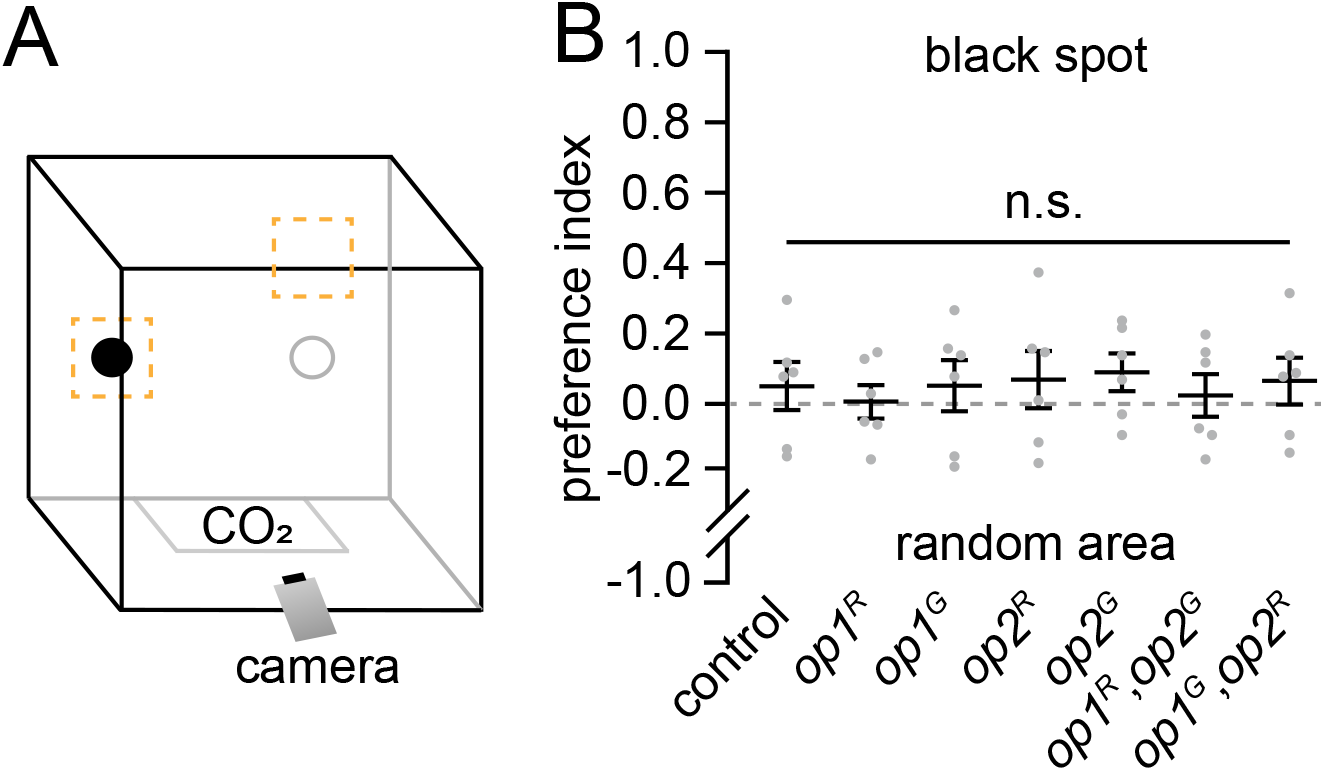
Testing preferences for the black spot versus a random area in a cage with standard air. (A) Schematic of the cage for performing the vision-guided host-seeking assay. The orange squares represent the regions used to document the trajectories around the black circle and a random area. (B) Vision-guided target selection cage assay for testing preferences of the indicated female mosquitoes for the black spot versus a random area under clean air conditions. n=6 (50 females/assay). Total trajectories: control, n=23—51; *op1^R^*, n=35—64; *op1^G^*, n=17—86; *op2^R^*, n=29—51; *op2^G^*, n=30—64; *op1^R^,op2^G^*, n=24—48; *op1^G^,op2^R^*, n=19— 42.

**Figure S3.**
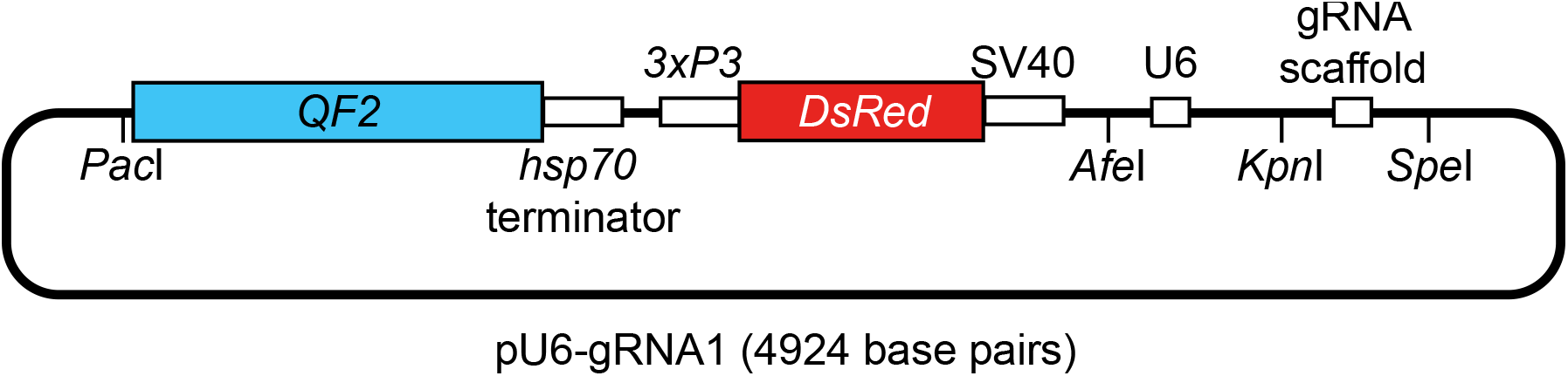
Schematic of the pU6-gRNA1 plasmid. The vector backbone contained the sequences encoding QF2 linked to the *hsp70* transcriptional terminator, DsRed driven by the *3XP3* promotor, and flanked on the 3’ end by the SV40 transcriptional terminator, the *U6* promotor, and gRNA scaffold. The following restriction endonuclease sites are indicated: *Pac*I for introducing the upstream homology arm, *AfeI* for introducing the downstream homology arm, and *KpnI* and *SpeI* for introducing the sgRNAs.

## Supplementary Tables

**Table S1.**
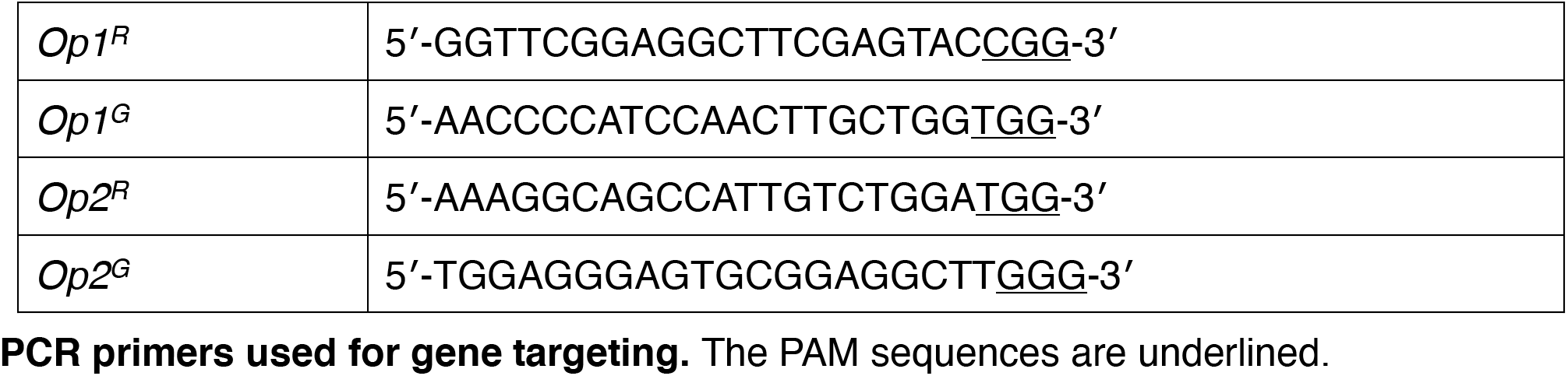
PCR primers used for gene targeting. The PAM sequences are underlined.

**Table S2.**
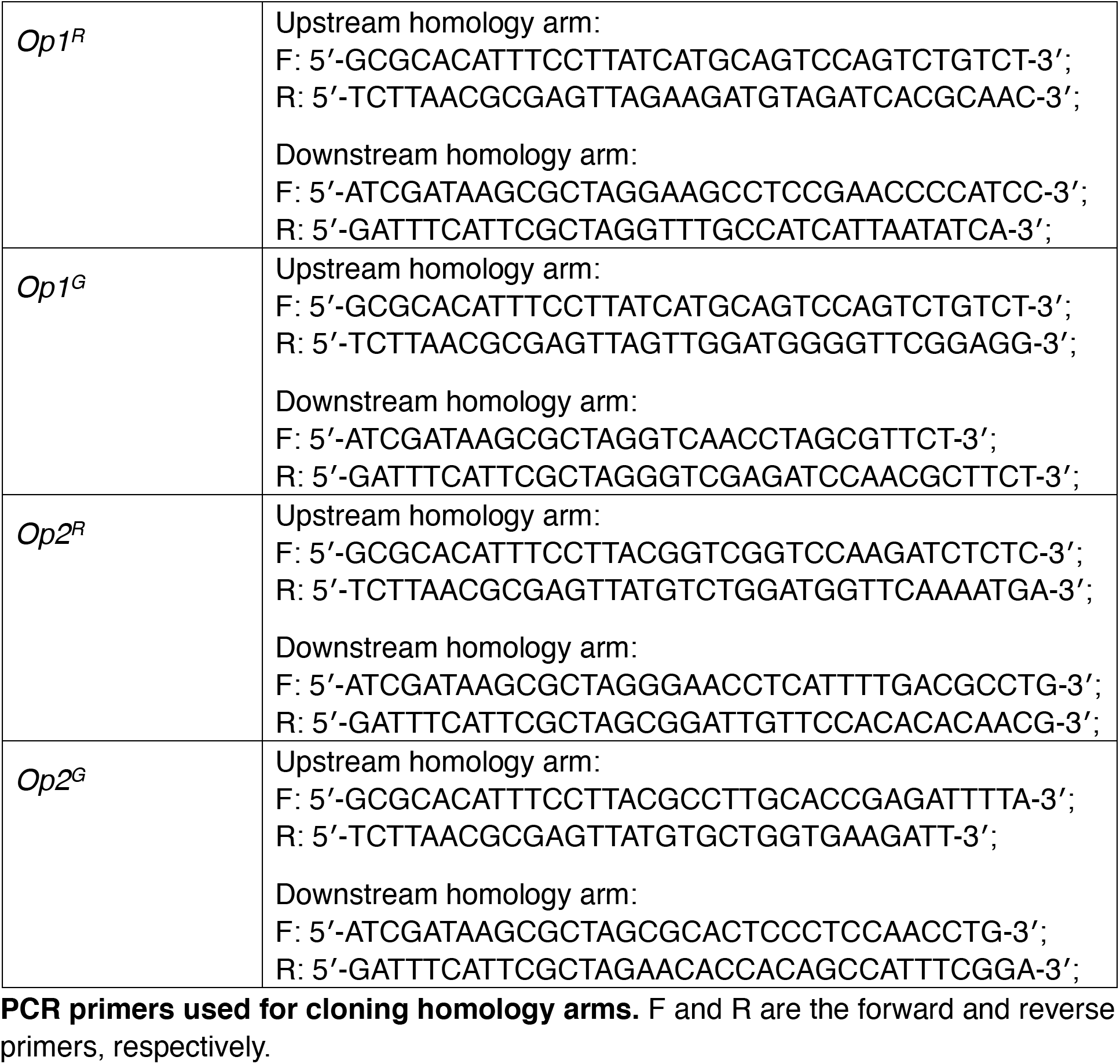
PCR primers used for cloning homology arms. F and R are the forward and reverse primers, respectively.

**Table S3.**
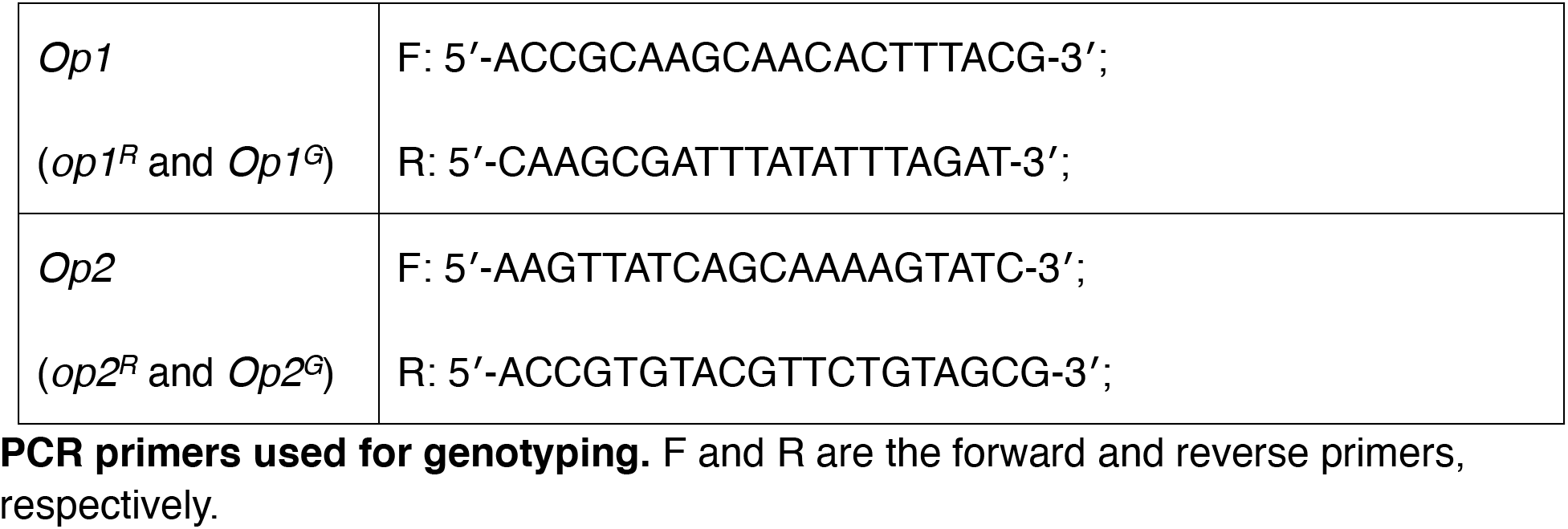
PCR primers used for genotyping. F and R are the forward and reverse primers, respectively.

**Table S4.**
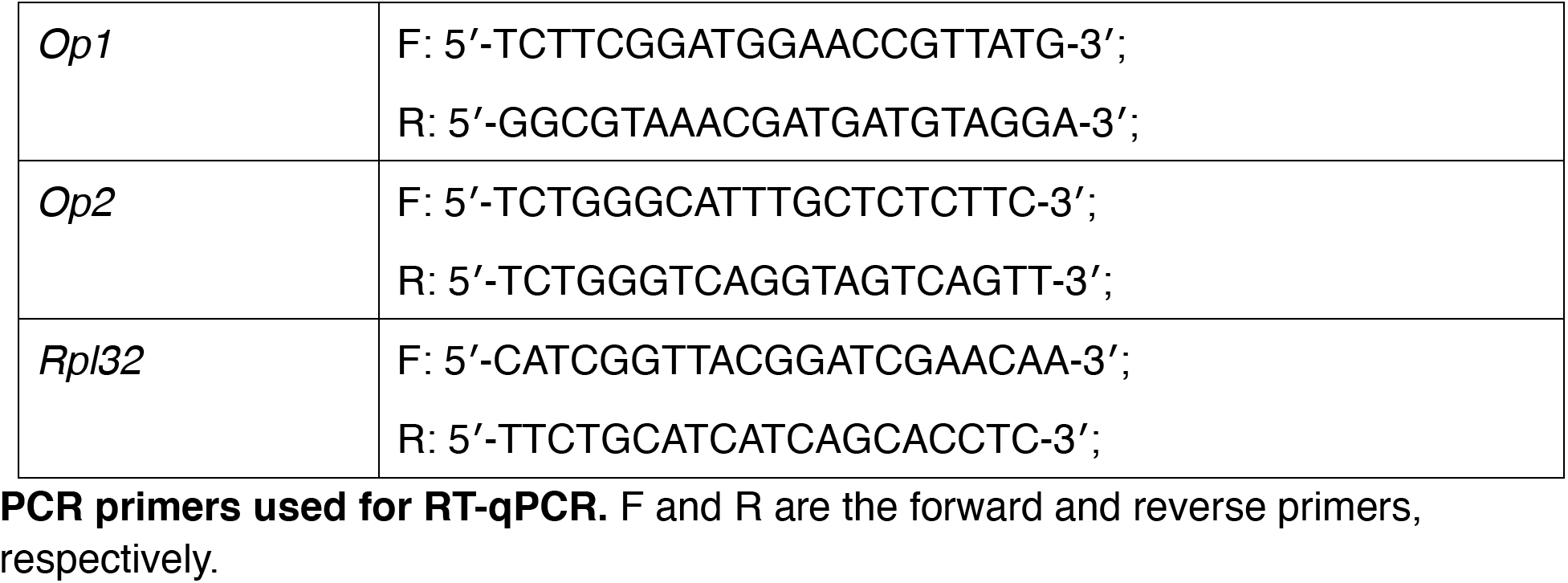
PCR primers used for RT-qPCR. F and R are the forward and reverse primers, respectively.

## Notes

### Competing Interest Statement

The authors have declared no competing interest.

